# The nervous system leverages the non-linear properties of the Achilles tendon to regulate ankle impedance during postural control

**DOI:** 10.1101/2022.07.27.501735

**Authors:** Kristen L. Jakubowski, Daniel Ludvig, Eric J. Perrault, Sabrina S.M. Lee

## Abstract

Regulating ankle impedance is essential for controlled interactions with the environment and rejecting unexpected disturbances. Ankle impedance in the sagittal plane depends strongly on the triceps surae and Achilles tendon, but their relative contributions remain unknown. It is commonly assumed that ankle impedance is controlled by changing muscle activation and, thereby, muscle impedance, but this ignores the fact that tendon impedance also changes with loading caused by increases in muscle activation. Thus, we sought to determine the relative contributions from the triceps surae and Achilles tendon during conditions relevant to postural control. We used a novel technique that combines B-mode ultrasound imaging with joint-level perturbations to quantify ankle, muscle, and tendon impedance simultaneously across activation levels from 0 – 30% of maximum voluntary contraction. We found that muscle and tendon stiffness, the static component of impedance, increased with voluntary plantarflexion contractions, but that muscle stiffness exceeded tendon stiffness at very low loads (21 ± 7 N). Above these loads, corresponding to 1.3% of maximal strength for an average participant in our study, ankle stiffness was determined predominately by Achilles tendon stiffness. Hence, the nervous system leverages the non-linear properties of the Achilles tendon to increase ankle stiffness during postural conditions.

## BACKGROUND

The ability to adapt the mechanical properties of the ankle is essential for seamlessly transitioning across different terrains when walking and for maintaining postural stability when unexpectedly perturbed [1, 2]. The triceps surae muscles and the Achilles tendon are the primary determinants of ankle mechanics in the sagittal plane, but their relative contributions remain largely unknown. It is commonly assumed that changes in ankle mechanics during active contractions are largely determined by the activation-dependent properties of muscle [3-5], but there have been limited *in vivo* measurements validating this presumption. Determining how the triceps surae and Achilles tendon mechanics contribute to ankle mechanics across a broad range of physiological conditions would provide fundamental insight into the mechanisms underlying humans’ ability to navigate their physical world. Such knowledge could also aid in developing targeted interventions when musculotendon mechanics are altered due to neuromuscular pathologies, or biomimetic assistive devices [4, 6]. As such, we sought to determine the relative contribution from the triceps surae and Achilles tendon to the mechanics of the ankle.

The assumed primary role of muscle in determining the mechanical properties of the ankle is based on two assumptions. The first is that muscle impedance is substantially lower than tendon impedance for most physiological conditions. Impedance—a quantitative measure of mechanics— describes the dynamic relationship between an imposed displacement and the evoked forces or torques [7]. Due to the serial connection between the muscle and tendon, ankle impedance will be determined mainly by the component with the lowest impedance when the impedance of each component differs substantially. The Achilles tendon is long and compliant [8, 9], and its impedance relative to that of the triceps surae is unknown. Therefore, muscle impedance may not be substantially lower than tendon impedance during physiologically relevant conditions.

The second assumption is that tendon impedance is constant across loads and that changes in joint impedance must therefore be due to changes in muscle impedance. Nearly all experimental studies quantifying muscle and tendon mechanics have focused on stiffness, the static component of impedance. It is well known from *in vivo* experiments that muscle stiffness changes with the activation-dependent changes in muscle force [10, 11]. Several studies have measured tendon stiffness *in vivo*, but often relying on the assumption that it remains constant across loads [12-15]. While this is a reasonable assumption at high loads (above approximately 30% of maximum force) [16, 17], tendons have a non-linear stress-strain relationship that results in load-dependent stiffness properties in the lower load regime within the “toe-region” [16, 17]. Ultimately, accounting for the non-linear properties of the tendon could impact the relative contributions from the muscle and tendon to the stiffness of the joint.

There is conflicting experimental evidence on how triceps surae and Achilles tendon stiffness vary with respect to each other and their relative contributions to ankle stiffness. This stems from the fact that few studies have examined muscle and tendon stiffness over a wide range of loads relevant to common functional tasks. Previously, it has been observed that tendon stiffness is greater than muscle in experiments that only considered activation levels above approximately 30% of the maximum voluntary contraction (MVC) [12]. In contrast, others have observed that the Achilles tendon is more compliant than the triceps surae during standing [18], which typically occurs around 15% MVC [19]. These conflicting results may be due partly to differences in the tested range of muscle activations. To our knowledge, no one has bridged the gap between these estimates and quantified the relative contribution from the muscle and tendon across a range of activations that are relevant to many functional tasks.

The objective of this study was to determine how the triceps surae and Achilles tendon contribute to the impedance of the ankle during conditions relevant to postural control. We used an innovative technique that combines joint-level perturbations with B-mode ultrasound to quantify ankle, muscle, and tendon impedance [20]. Given the limited and conflicting experimental data reported in the literature, we tested the null hypothesis that the muscle and tendon contribute equally to ankle impedance to determine which structure was most dominant over contraction levels ranging from 0 to 30% MVC. Our results help determine the mechanisms contributing to the regulation of human ankle impedance, as needed for seamless interactions with the environment. As a secondary objective, we quantified the frequency ranges over which muscles and tendons behave elastically. Though there are conditions for which muscles and tendons exhibit spring-like behavior, both structures are viscoelastic [21, 22]. Until recently [20], it has not been possible to quantify muscle and tendon impedance *in vivo* in humans. As such, it is unknown under which conditions it is reasonable to assume that muscles and tendons behave as simple springs with only a stiffness component and when they exhibit more complex mechanical properties. We, therefore, quantified these regimes in this study.

## METHODS

### Participants

Seventeen healthy young adults (age = 27 ± 3 years (mean ± standard deviation); height = 1.7 ± 0.1 m; body mass = 73 ± 15 kg, 8 males and 9 females) participated in this experiment. All participants were right leg dominant and had no history of neuromuscular or musculoskeletal injuries to their right leg. All participants provided informed consent prior to participation. The Northwestern University Institutional Review Board approved the study, and all methods were carried out according to the approved protocols (STU00009204 & STU00213839).

### Experimental setup

Participants were seated in an adjustable chair (Biodex Medical Systems, Inc. Shirley, NY), with their trunk and torso stabilized with safety straps (Fig 1). Participants’ right leg was extended in front of them with their knee flexed at 15°. A knee brace (Innovator DLX, Ossur, Reykjavik, Iceland) stabilized the knee in this position. The participant’s right foot was attached rigidly to an electric rotary motor (BSM90N-3150AF, Baldor, Fort Smith, AR) via a custom-made fiberglass cast at an ankle angle of 90°. The cast encased the entire foot, extending distally from the medial and lateral malleoli to the toes, thus preserving the full range-of-motion of the ankle but preventing any movement of the foot or toes. The axis of rotation of the motor was aligned with the ankle center of rotation in the sagittal plane, restricting all movement to the plantarflexion/dorsiflexion direction. Electrical and mechanical safety stops limited the rotation of the motor within the participant’s range of motion. A 24-bit quadrature encoder integrated with the motor measured ankle angle (24-bit, PCI-QUAD04, Measurement Computing, Norton, MA), while a 6-degree-of-freedom load cell (45E15A4, JR3, Woodland, CA) measured all ankle forces and torques. Throughout the experiment, the motor was controlled in real-time via xPC target (MATLAB, Mathworks, Natick, MA).

**Figure 1.**
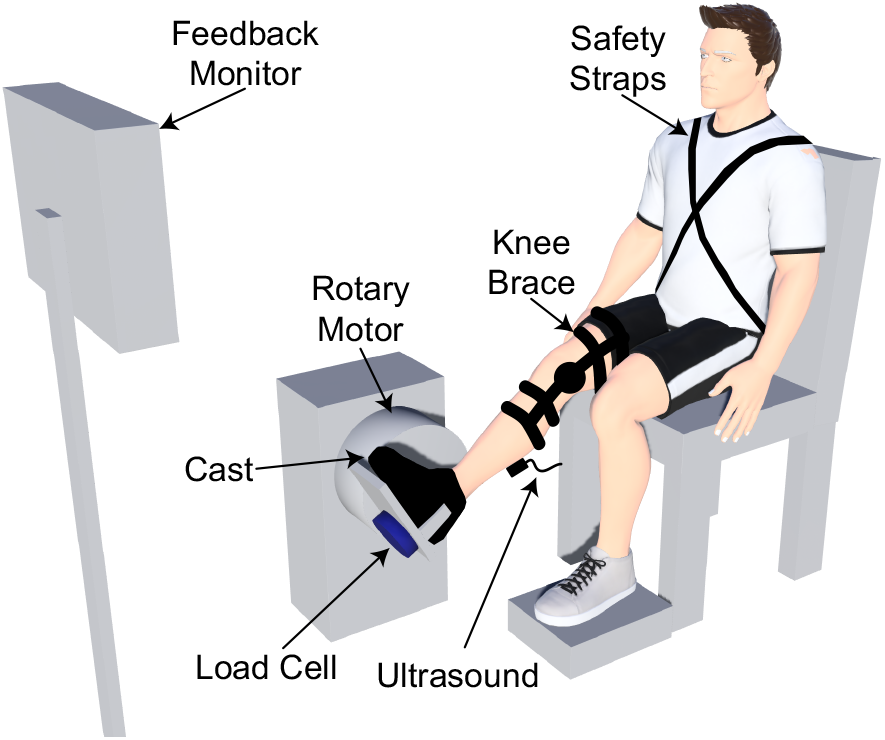
Schematic of the experimental setup. A custom-made cast secured the subject’s foot to the rotary motor. The rotary motor rigidly controlled the ankle joint angle, while the load cell measured the resultant ankle torque. We used B-mode ultrasound to image the muscle-tendon junction of the medial gastrocnemius. The knee brace secured the knee in a stable position, preventing unwanted knee flexion or extension. The feedback monitor provided real-time feedback on the magnitude of the plantarflexion torque and the tibialis anterior muscle activity.

Single differential bipolar surface electrodes (Bagnoli, Delsys Inc, Boston, MA) measured muscle activity from the medial and lateral gastrocnemius and soleus (ankle plantarflexors) and the tibialis anterior (ankle dorsiflexor). Standard skin preparation methods were performed prior to electrode placement [23], and electrodes were placed on the belly of the muscle. Electromyography (EMG) signals were amplified to maximize the signal resolution in each channel. EMG data were collected for visual feedback provided to the subjects. All analog data were passed through an antialiasing filter (500 Hz using a 5-pole Bessel filter) and sampled at 2.5 kHz (PCI-DAS1602/16, Measurement Computing, Norton, MA, USA).

A B-mode ultrasound system using a linear transducer (LV7.5/60/128Z-2, LS128, CExt, Telemed, Lithuania) recorded images of the medial gastrocnemius muscle-tendon junction (MTJ). A custom-made probe holder and elastic adhesive wrap (Coban™, 3M, St. Paul, MN) secured the probe to the leg. We positioned the ultrasound probe to center the MTJ on the image. At the start of ultrasound data collection, a trigger signal was used to synchronize the ultrasound data collection with all other measurements. Ultrasound images were acquired with a mean frame rate of 124 Hz. All ultrasound data were saved for processing offline.

### Protocol

At the start of each experiment, we collected three 10-second isometric maximum voluntary contractions (MVC) trials in both the plantarflexion and dorsiflexion directions. These data were used to scale the visual feedback provided to the participants.

Our primary objective was to determine how muscle, tendon, and ankle impedance vary across various levels of plantarflexion torque. This was accomplished by instructing participants to produce different levels of isometric plantarflexion torque while the rotary motor applied small rotational perturbations in the sagittal plane. We used pseudo-random binary sequence (PRBS) perturbations with an amplitude of 0.175 radians, a maximum velocity of 1.75 radians per second, and a switching time of 153 ms. We tested seven isometric plantarflexion torque levels from 0% to 30% MVC in 5% increments. Participants were provided real-time visual feedback of their normalized plantarflexion torque. Tibialis anterior EMG was also provided to prevent co-contraction. Rectified EMG and torque signals were low pass filtered at 1 Hz to remove high-frequency components from the applied perturbations (2^nd^ order Butterworth). Subjects completed three trials at each level of plantarflexion torque in a randomized fashion. Each trial lasted 65 seconds. Rest breaks were provided as needed between trials to prevent fatigue.

The measured ankle torque included the gravitational and inertial contributions from the apparatus connecting the foot to the motor. A single trial was collected with only the cast attached to the rotary motor enabling us to remove these contributions from the net torque measured in each trial.

### Data processing and analysis

All data were processed and analyzed using custom-written software in MATLAB. The same individual manually digitized the MTJ within each frame of the ultrasound videos [20]. All ultrasound data were resampled using linear interpolation to match the sampling rate of the other experimental signals (2.5 kHz).

We computed ankle, muscle, and tendon impedance as described previously [20]. Briefly, we used non-parametric system identification to estimate ankle, muscle, and tendon impedance from the experimental measures of ankle angle, ankle torque, and displacement of the MTJ (Fig 2). We quantified ankle impedance as the relationship between the imposed ankle rotations and the resultant ankle torque [7]. Measurement of the MTJ motion allowed us to estimate muscle and tendon impedance under the assumption that the muscle and tendon are connected in series [24], and that the displacement of the muscle-tendon unit is determined by the angular rotation of the ankle multiplied by the Achilles tendon moment arm. We refer to the relationship between MTJ displacement and the angular rotations of the ankle as the translation ratio. Specifically, to characterize ankle, muscle, and tendon impedance, we estimated ankle impedance and the translation ratio, and used these quantities to compute muscle and tendon impedance [20]. We previously demonstrated that the magnitude of the frequency response functions was nearly constant from 1 to 3 Hz, indicating that stiffness was the dominant contributor to impedance over this frequency range [20]. As such, we computed the stiffness component of ankle, muscle, and tendon impedance by averaging the magnitude of the respective frequency response functions from 1 to 3 Hz. Our primary analysis will focus on the stiffness component of impedance due to its relevance in the control of posture and movement at the ankle [25].

**Figure 2.**
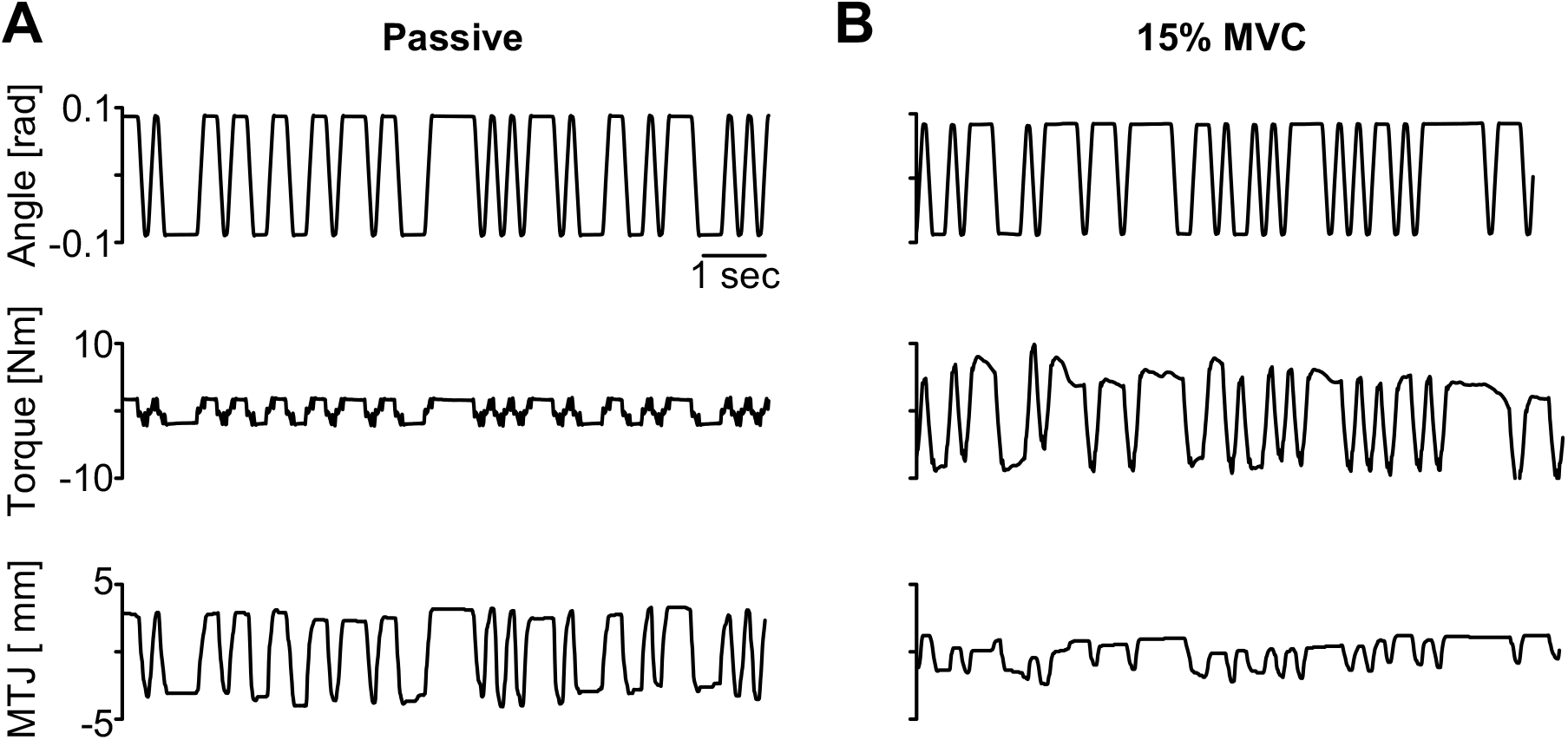
Representative data used to estimate ankle, muscle, and tendon impedance. Representative data from a passive trial (A) and a trial when the participant was instructed to maintain 15% of their maximum voluntary torque (MVC) (B). The rotary motor rigidly controlled the position of the participant’s ankle (angle) at all times. We measured the resultant ankle torque and muscle-tendon junction (MTJ) displacement from the medial gastrocnemius resulting from the applied random perturbations. Torque and MTJ displacement have been detrended.

A single approximation of the Achilles tendon moment arm (51.4 mm) was used for all analyses. This was estimated as the mean across subjects from Clarke et al. [26] with an ankle angle of 90°. It has been demonstrated that the Achilles tendon moment arm does not scale with anthropometric data [26, 27]. Additionally, system identification is a quasi-linear approximation about a single operating point, which, in our study, was 90°. Therefore, we approximated the moment arm as a single value.

Ankle and tendon stiffness varied non-linearly with plantarflexion torque (or musculotendon force). Therefore, the ankle and tendon stiffness experimental data were fit with non-linear models to synthesize our results. The model used to characterize torque-dependent changes in ankle stiffness was:

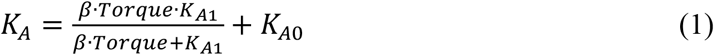

in which *K*_*A*_ represents the modeled ankle stiffness, *Torque* was the input to the model, *β, K*_*A1*_ and *K*_*A0*_ are the optimized parameters. A similar model has been used to characterize load-dependent changes in the stiffness of a muscle-tendon unit [28].

Tendon stiffness was modeled by an exponential function:

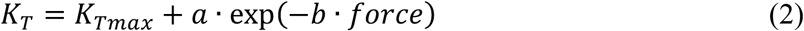

in which *K*_*T*_ represents the modeled tendon stiffness, musculotendon *force* was the input to the model, and *K*_*Tmax*_, *a*, and *b* are the optimized parameters. This model was chosen since exponential models have been used previously to characterize the non-linear toe-region of the tendon stress-strain curve [29]. We computed musculotendon force by dividing the measured ankle torque by the Achilles tendon moment arm.

*Sensitivity analyses*

We evaluated the sensitivity of ankle stiffness to changes in muscle and tendon stiffness at different levels of force. We first consider that ankle stiffness (*K*_*A*_) is determined by the serial connection of the muscle and tendon and can be described as a function of these stiffnesses [20], such that:

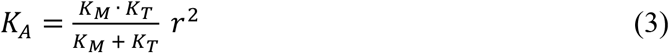

where *r* represents the Achilles tendon moment arm in the sagittal plane, *K*_*M*_ represents muscle stiffness, and *K*_*T*_ represents tendon stiffness. This relationship was used to derive the sensitivity of ankle stiffness to muscle and tendon stiffness using Eq 4, where *S*_*x*_ is the relative sensitivity to a given parameter *x* (either muscle or tendon stiffness).

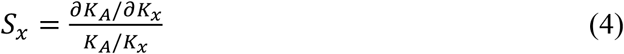

The average values of muscle and tendon stiffness estimated from our experiment were used to compute numerical values for the sensitivity of ankle stiffness.

### Statistical analysis

We sought to determine how the triceps surae and Achilles tendon contribute to the impedance of the ankle over a range of activation levels. Non-linear mixed-effects models were used to characterize the ankle stiffness–torque relationship and the tendon stiffness–force relationship (Eq 1 & 2). A linear mixed-effects model was used to describe the muscle stiffness–musculotendon force relationship. For all models, subject was treated as a random factor, and plantarflexion torque or musculotendon force was a continuous factor. A restricted maximum likelihood method was used to estimate all models [30]. The model fit for ankle, muscle, and tendon stiffness was assessed by quantifying the coefficient of determination (R^2^) for each participant from the respective mixed-effects model. We tested the null hypothesis that the muscle and tendon contribute equally to ankle stiffness. We used a bootstrapping procedure to determine the range of musculotendon forces when muscle and tendon stiffness were not significantly different from each other to a level of p>0.05. The bootstrapping involved randomly resampling the data from each subject with replacement to create a new dataset for the entire pool of subjects. This process was repeated 200 times. Each synthesized dataset was analyzed as described above to create a distribution of estimates for which muscle and tendon stiffnesses were the same. Our null hypothesis—that muscle and tendon stiffness contribute equally to the stiffness of the ankle—was accepted within the 95% confidence intervals of this distribution and rejected elsewhere. All metrics reported are mean ± 95% confidence intervals unless otherwise noted.

## RESULTS

### Muscle stiffness exceeded tendon stiffness at low loads

At all levels of activation, the magnitudes of the frequency response functions for muscle and tendon impedance were nearly constant from 1 – 6.5 Hz (Fig 3), indicating that stiffness is the primary contributor to impedance at these frequencies. Therefore, it is reasonable to assume that muscle and tendon behave as simple springs during the conditions tested and over this frequency range, which is not impacted by changes in load.

**Figure 3.**
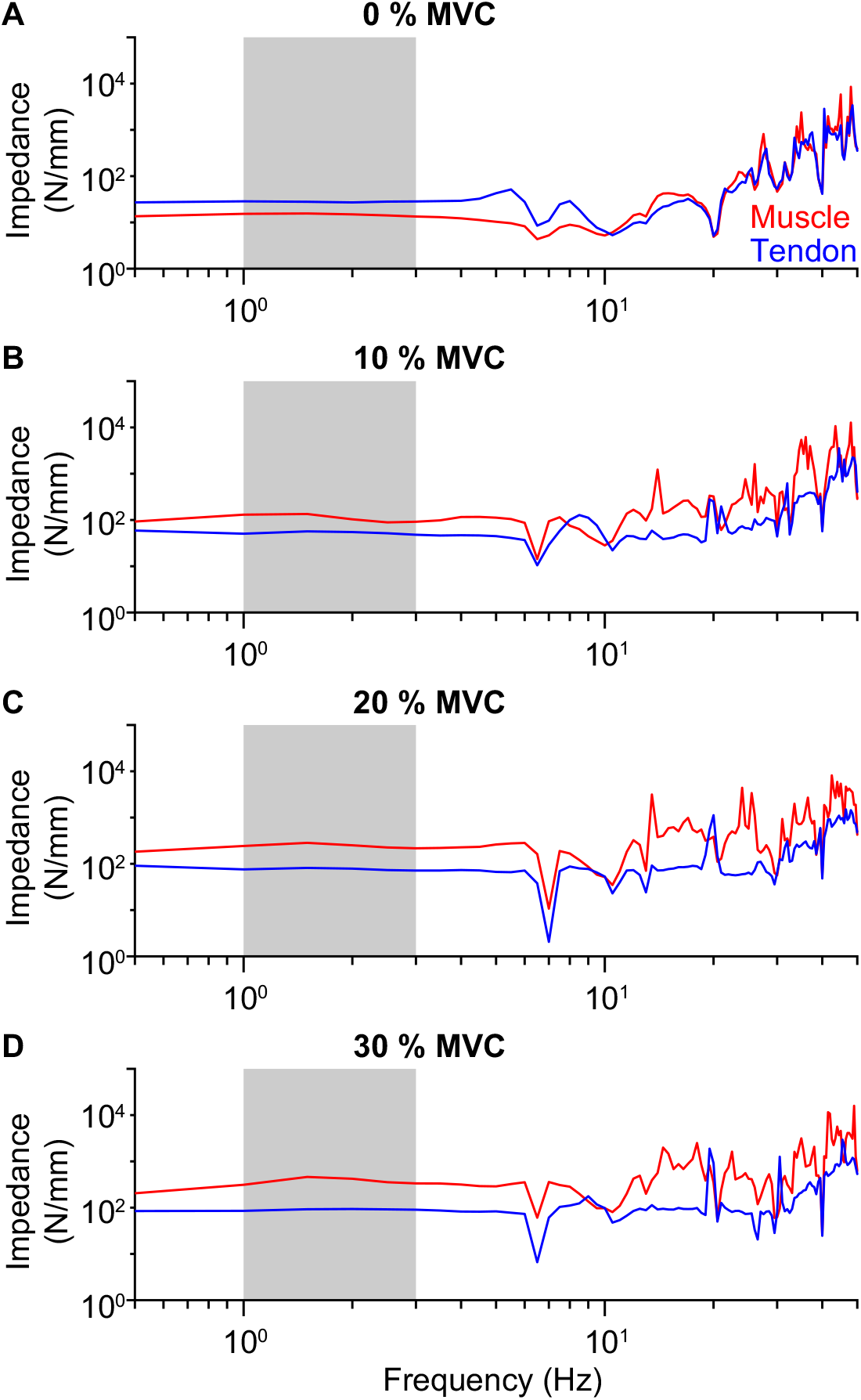
Stiffness is the dominant contributor to muscle and tendon impedance at low frequencies. Muscle (red) and tendon (blue) impedance frequency response functions from a representative participant at (A) 0% MVC, (B) 10% MVC, (C) 20% MVC, and (D) 30% MVC. The magnitudes of the frequency response functions were nearly constant from 1 – 6.5 Hz, indicating that stiffness is the primary contributor to impedance. Values between 1 – 3 Hz were used to compute stiffness (shaded region).

Muscle and tendon stiffness increased with increases in musculotendon force (Fig 4). Figure 4A displays the experimental measures and model fits from an individual subject. The muscle and tendon stiffness models fit the data well for the representative subject (muscle: R^2^ = 0.94; tendon: R^2^ = 0.91), and across the entire group (muscle: R^2^ = 0.94 ± 0.01; tendon: R^2^ = 0.94 ± 0.01).

**Figure 4.**
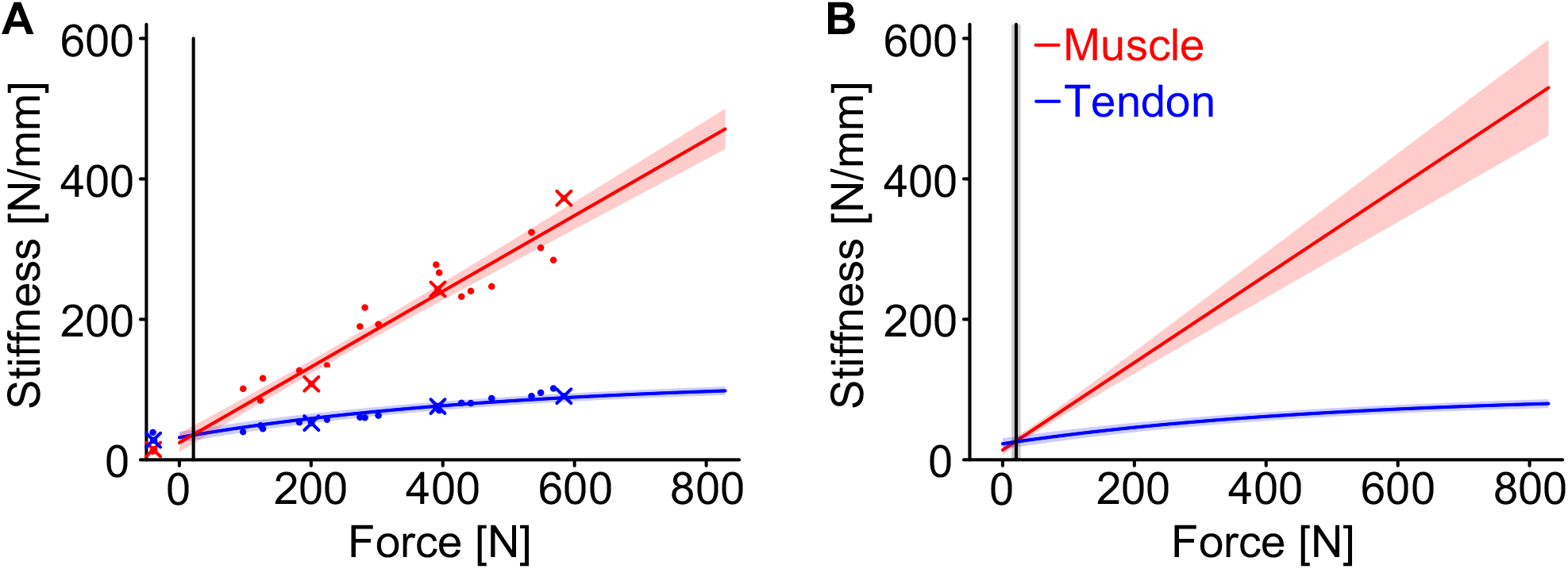
Muscle stiffness exceeded tendon stiffness past the lowest levels of force. (A) Muscle stiffness (red) and tendon stiffness (blue) for an individual subject, illustrating that muscle stiffness exceeded tendon stiffness at low levels of musculotendon force (21 N – solid black line). Each point represents an individual trial. The x’s correspond to the trials illustrated in Fig 3. (B) This trend was preserved across all subjects (n=17). Muscle stiffness exceeded tendon stiffness at 21 ± 7 N (mean ± 95% CI – solid line and shaded area). The tendon stiffness experimental data were modeled using Eq 2, while the muscle stiffness experimental data were modeled linearly. Mixed-effects models were used for muscle and tendon stiffness to account for random variability between subjects. The solid line indicates the estimated muscle and tendon stiffness from the respective mixed-effects models, with the shaded region being the 95% confidence intervals. We evaluated the range of musculotendon forces when muscle and tendon stiffness were not significantly different from each other to a level of p > 0.05 using a bootstrapping procedure. The solid black line was the mean musculotendon force where muscle and tendon stiffness were equivalent within the set level of statistical significance, with shading indicating the 95% confidence intervals across the bootstrapped samples.

We found that muscle stiffness increased at a greater rate with increases in force than tendon stiffness. A representative participant shown in Fig 4A, illustrates that muscle stiffness was greater than tendon stiffness at 21 N (solid line). This trend was consistent across all subjects. We observed that muscle stiffness exceeded tendon stiffness at 21 ± 7 N (Fig 4B). The musculotendon force where muscle stiffness exceeded tendon stiffness (21 N) occurred at a very low contraction level, corresponding to 1.5 ± 0.2% of the maximum voluntary torque across all subjects. At the highest force tested in this study, ∼830 N, the muscle was approximately 6.6 times stiffer than the tendon.

### Ankle stiffness was most sensitive to changes in tendon stiffness

A unique feature of our measurement technique is that we were able to quantify ankle, muscle, and tendon stiffness simultaneously, enabling us to quantify the relative contributions from the muscle and tendon to the stiffness of the joint. As others have reported [3, 7, 31], we found that ankle stiffness increased with voluntary contraction (Fig. 5). This increase was non-linear and described well by Eq. 1 for individual subjects (Fig. 5A; R^2^ = 0.99), and the full population of tested subjects (Fig. 5B; R^2^ = 0.98 ± 0.007). Our values of ankle stiffness are consistent with previous reports using a similar experimental protocol [32].

**Figure 5.**
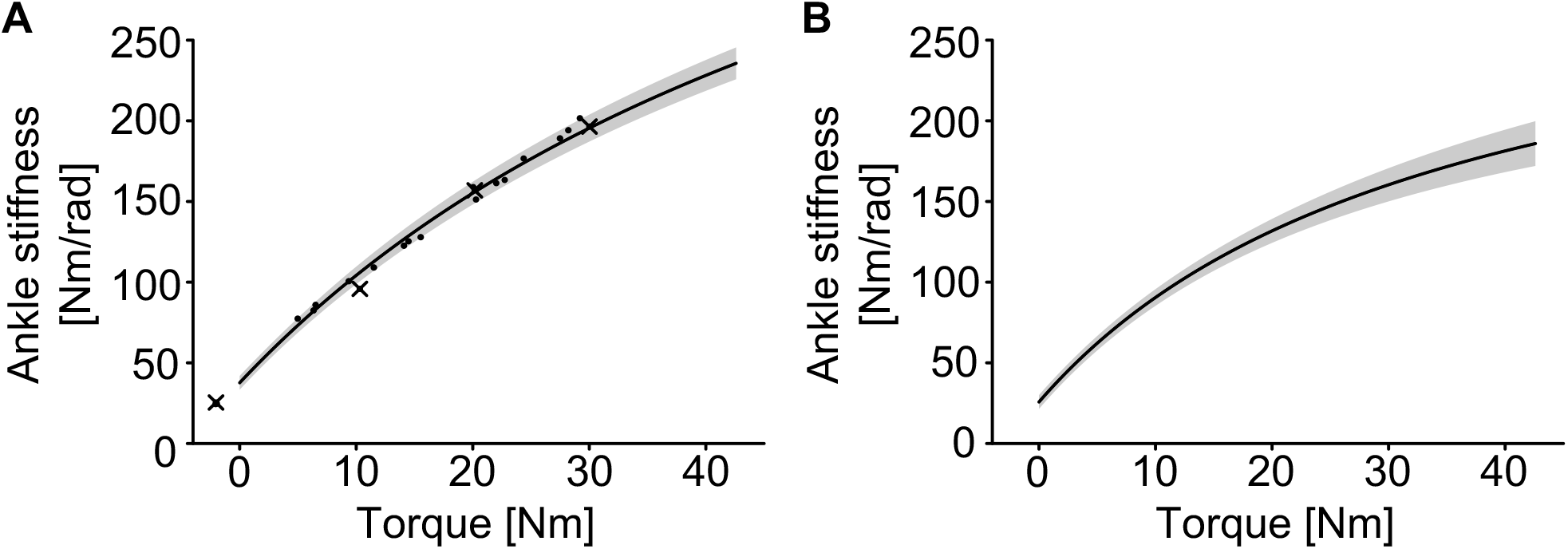
Ankle stiffness increased with increased plantarflexion torque. A) Ankle stiffness estimates for an individual subject, illustrating the increase in stiffness with torque. Each point represents an individual trial. The x’s correspond to the trials illustrated in Fig 3. This trend was preserved in the group results (n=17) (B). The ankle stiffness experimental data were modeled using Eq 1. A mixed-effects model was used to account for random variability between subjects. For all plots, the solid line indicates the estimated stiffness from the respective fitted model, with the shaded region being the 95% confidence intervals.

We completed a sensitivity analysis to quantify how changes in muscle and tendon stiffness influence ankle stiffness across the range of tested forces (Fig 6). As expected, ankle stiffness was most sensitive to the tendon for forces above 21 N, where tendon stiffness became lower than muscle stiffness. For forces above 350 N, corresponding to approximately 20% MVC in our population of subjects, ankle stiffness was nearly 4 times more sensitive to changes in tendon stiffness than to changes in muscle stiffness. The importance of tendon stiffness for determining ankle stiffness increased at further contraction levels. These results provide additional evidence that the mechanical properties of the human ankle are determined primarily by the non-linear mechanical properties of the Achilles tendon.

**Figure 6.**
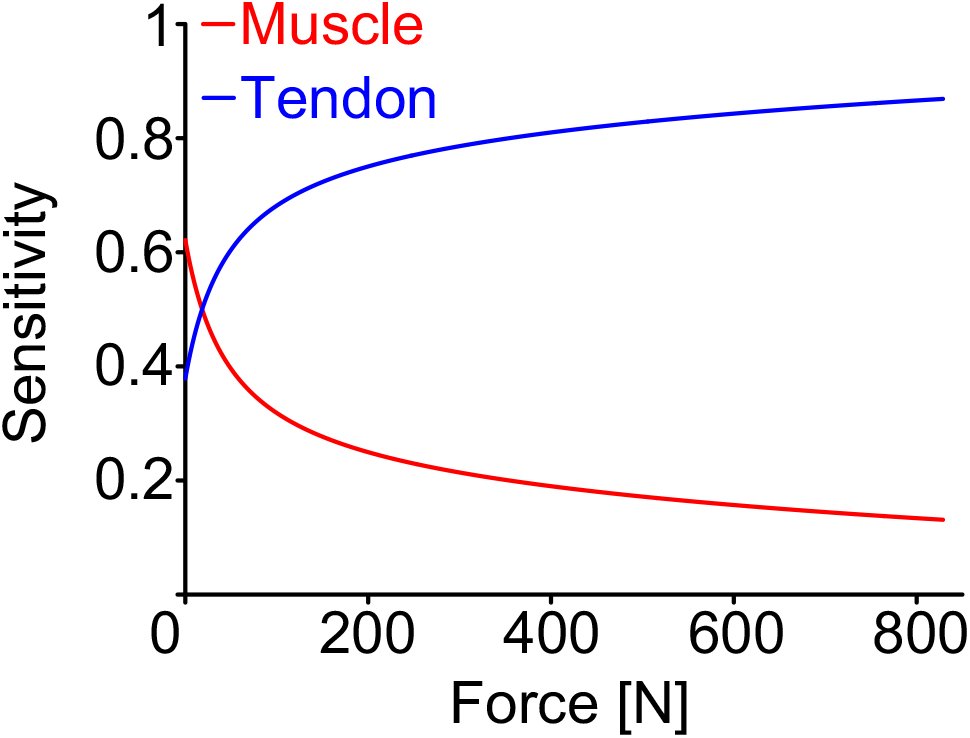
Sensitivity of ankle stiffness to changes in muscle (red) and tendon (blue) stiffness. Beyond the lowest levels of force, ankle stiffness was more sensitive to changes in tendon stiffness compared to changes in muscle stiffness.

## DISCUSSION

Regulating ankle impedance is critical when adapting to varying environmental conditions and responding to postural disturbances. This study sought to determine how the triceps surae and Achilles tendon contribute to the sagittal plane impedance of the ankle over a range of activation levels. We used our novel technique to quantify ankle, muscle, and tendon impedance simultaneously [20]. We found that both muscle and tendon impedance increased with activation, and that both had spring-like properties for frequencies below approximately 6.5 Hz. Muscle stiffness exceeded tendon stiffness beyond the lowest forces and levels of activation (∼21 N or ∼1.5 ± 0.2% MVC). The stiffness of the human ankle during plantarflexion is determined largely by the net stiffness of the serially connected Achilles tendon and triceps surae muscles. Because springs connected in series have a net stiffness that is limited by the most compliant (least stiff) element, our results indicate that the mechanical properties of the Achilles tendon, a passive structure, have a substantial impact on the activation-dependent increases in ankle stiffness at almost all levels of muscle activation. This finding is in contrast to the common assumption that the regulation of ankle stiffness is directly linked to activation-dependent changes in muscle stiffness [3-5]. Instead, our results suggest that the nervous system leverages the non-linear properties of the Achilles tendon to increase ankle stiffness during postural conditions. This is a fundamental shift in the assumed roles of the muscle and tendon and our current understanding of how muscle and tendon impedance contribute to the impedance of the ankle.

### Characteristics of triceps surae and Achilles tendon impedance

We found that muscle and tendon have spring-like properties below approximately 6.5 Hz, as indicated by the nearly constant-valued impedance frequency response functions (Fig 3). This result implies that viscous and inertial properties of the muscle and tendon are small relative to stiffness over this frequency range. This result is consistent with testing in excised tendons, where it has been found that the estimated tendon mechanical properties are invariant with respect to the frequency of the applied stretch up to 11 Hz [33, 34]. It is also consistent with previous findings within feline muscle, where the muscle behaves elastically in response to small stochastic perturbations over a similar frequency range to what we tested [35]. Our measured response in muscle suggests that our measurements remained within its short-range stiffness region [10]. Muscle short-range stiffness describes the initial response to small, fast displacements prior to reflexive or volitional muscle activation and is critical in the control of posture and limb stability [35, 36]. This finding is consistent with our previous results that demonstrated that our muscle stiffness estimates are similar to measurements of muscle short-range stiffness scaled to the triceps surae [20]. We do note, however, that if the stretch within the muscle or tendon was larger or slower, we would expect to observe more complex viscoelastic behavior. For example, when a larger stretch is applied to a muscle, the response is no longer purely elastic [10]. Similarly, within tendon, when stretch velocity is slower, the mechanical properties of the tendon decrease [37].

Our estimated values of muscle stiffness were larger than the few previous reports that attempted to quantify the stiffness of the human triceps surae muscle *in vivo*. This is likely due to the small size of our perturbations compared to earlier studies. All previous estimates of human triceps surae muscle stiffness used perturbations at least twice as large as those we applied (20° or larger) [12, 38]. Previously, Hauraix et al. [12] reported a triceps surae muscle stiffness value of 218 N/mm at 40% MVC, while Clark et al. [38] reported a muscle stiffness of 118 N/mm at 25% MVC. For comparison, we estimate muscle stiffness to be 261 N/mm at 25% MVC for an average participant in our study. Muscle stiffness varies based on the size of the applied perturbation [10]. Therefore, given the difference in perturbation size, it was expected that the previously reported muscle stiffness values would be lower than our results. Our novel *in vivo* estimates of muscle stiffness may be especially pertinent for stability and the response to unexpected postural disturbances when the short-range stiffness of the muscle is important.

The observed increase in tendon stiffness with increases in musculotendon force suggests that the Achilles tendon was within the non-linear toe-region of its stress-strain curve during our experiments (Fig 4). Tendons exhibit a strain-dependent increase in stiffness at low strains (e.g., the toe-region of the stress-strain curve) [16]. While Achilles tendon stiffness has been characterized before [12-15], previous studies have only estimated its stiffness above 30% MVC to satisfy the methodological assumption that tendon stiffness is constant. Our approach is not constrained by this assumption, allowing measurements to be made at lower forces corresponding to activation levels that occur during everyday activities like standing and walking [19].

### Limitations

Our technique for estimating muscle and tendon stiffness assumes that all plantarflexion torque is transmitted through the Achilles tendon to the triceps surae, omitting contributions from other structures that span the joint (e.g., the joint capsule and other musculotendon units) [20]. This assumption is valid during plantarflexion contractions, when the musculotendon force from the triceps surae is significantly greater than contributions from other sources. However, other structures can have a substantial effect relative to Achilles tendon force when the ankle is passively dorsiflexed. To mitigate their contributions, we positioned the ankle in a neutral position where passive torque is minimal [39]. We may still be overestimating muscle and tendon stiffness during passive conditions, but this limitation will have a negligible impact on our main conclusions when the triceps surae are active.

### Functional implications and Conclusions

While the data presented was during isometric conditions, our findings may explain an underlying physiological mechanism of previous estimates of ankle impedance during walking. Rouse et al. [40] observed that ankle stiffness estimated using perturbations of ankle posture during the stance phase of walking was similar to that estimated by the slope of the ankle torque – ankle angle relationship, also known as quasi-stiffness. This was surprising since these two estimation approaches can only yield the same results if the system is purely elastic and passive [41]. However, it is well documented that the triceps surae are active during the stance phase of locomotion [8, 40]. One possible explanation for the Rouse et al. [40] findings is that ankle stiffness was determined primarily by the Achilles tendon—a passive elastic structure—during the stance phase of walking, where it has been shown that muscle fascicle length changes are modest [8, 9], as in our postural experiment.

We observed that the Achilles tendon is less stiff than the triceps surae at almost all loads, but these results may not apply to other muscle-tendon units. For the Achilles tendon, the compliance of the tendon is essential for the storage and return of elastic energy, increasing the economy of locomotion [8, 9, 42, 43]. However, the mechanical properties of the muscle relative to the tendon will depend upon the functional role of each muscle-tendon unit and its corresponding architecture [44]. For example, muscles that have a similar fascicle length and tendon slack length have been termed “stiff”, while muscles where the fascicles are much shorter than the tendon—like the triceps surae—have been termed “compliant” [16]. It is almost certain that muscles in the former category will contribute more to the stiffness of the joint that they cross.

Finally, our results have implications for targeted rehabilitation. Changes in Achilles tendon stiffness that occur as a result of injury [45], or healthy aging [46] will impact ankle stiffness. For example, our results suggest that the previously reported age-related decrease in Achilles tendon stiffness will decrease the stiffness of the ankle for a fixed level of contraction [47]. This decrease could impair the control of posture and movement. To improve balance during tasks that require effective ankle stabilization, altering muscle stiffness through strength training might be less effective than increasing tendon stiffness through high magnitude loading [48, 49]. Ultimately, understanding the relative contributions from the muscle and tendon advances our fundamental understanding of how ankle stiffness is varied for an individual’s interactions with their physical world, and aids in developing targeted interventions when musculotendon mechanics are altered as a result of neuromuscular pathologies or aging.

